# Neuroepithelial progenitors generate and propagate non-neuronal action potentials across the spinal cord

**DOI:** 10.1101/2020.05.23.111955

**Authors:** Kalaimakan Hervé Arulkandarajah, Guillaume Osterstock, Agathe Lafont, Hervé Le Corronc, Nathalie Escalas, Silvia Corsini, Barbara Le Bras, Juliette Boeri, Antonny Czarnecki, Christine Mouffle, Erika Bullier, Elim Hong, Cathy Soula, Pascal Legendre, Jean-Marie Mangin

## Abstract

In the developing central nervous system, electrical signaling is thought to rely exclusively on differentiating neurons as they acquire the ability to generate action potentials. Accordingly, the neuroepithelial progenitors (NEPs) giving rise to all neurons and glial cells during development have been reported to remain electrically passive. Here, we investigated the physiological properties of NEPs in the mouse spinal cord at the onset of spontaneous neural activity (SNA) initiating motor behavior in embryos. Using patch-clamp recordings, we discovered that spinal NEPs exhibit spontaneous membrane depolarizations during episodes of SNA. These recurrent depolarizations exhibited a ventral-to-dorsal gradient with the highest amplitude located in the floor-plate – the ventral-most part of the neuroepithelium. Paired-recordings revealed that NEPs are extensively coupled via gap-junctions and form a single electrical syncytium. Although other NEPs were electrically passive, we discovered that floor-plate NEPs have the unique ability to generate large Na^+^/Ca^++^ action potentials. Unlike neurons, floor-plate action potentials relied primarily on the activation of voltage-gated T-type calcium channels (TTCCs). *In situ* hybridization showed that all 3 known subtypes of TTCCs are highly and predominantly expressed in the floor-plate. During SNA, we found that acetylcholine released by motoneurons recurrently trigger floor-plate action potentials by acting through nicotinic acetylcholine receptors. Finally, by expressing the genetically encoded calcium indicator GCaMP6f in the floor plate, we demonstrated that neuroepithelial action potentials are associated with calcium waves and propagate along the entire length of the spinal cord. By unraveling a novel physiological mechanism generating electrical signals which can propagate independently from neurons across a neural structure, our work significantly changes our understanding of the development, origin and extent of electrical signaling in the central nervous system.

**HIGHLIGHTS:**

- Spinal neuroepithelial progenitors (NEP) are depolarized during spontaneous neural activity
- NEPs form a single electrical syncytium connected by gap junctions
- Floor-plate NEPs generate large Na^+^/Ca^++^ action potentials in response to acetylcholine
- Neuroepithelial action potentials propagate across the entire spinal cord

## INTRODUCTION

Spontaneous neural activity (SNA) is a hallmark of developing neural networks and has been shown to regulate various developmental processes such as neuronal differentiation, migration, axon guidance and synapse formation (Blankenship and Feller, 2010; Kirischuk et al., 2017; Moody and Bosma, 2005; Spitzer, 2006). Until now, the electrical signals generated during SNA have been found to originate exclusively from differentiating neurons as they acquire the ability to fire action potentials and form their first functional synapses. By contrast, neuroepithelial progenitors (NEPs) – the stem cells of the central nervous system - have never been reported to actively generate or propagate any form of electrical signals, although they can be passively depolarized and/or exhibit calcium events in response to neurotransmitters released by developing neurons (Corlew et al., 2004; Kirischuk et al., 2017; LoTurco et al., 1995; Noctor et al., 2002; Rosa et al., 2015; Vitali et al., 2018; Wang et al., 2009; Weissman et al., 2004).

SNA was initially described in the embryonic spinal cord where it triggers the first motor rhythmic behavior (Hamburger and Balaban, 1963; Suzue and Shinoda, 1999), necessary for the proper development of the neuro- muscular and musculo-skeletal systems of vertebrates (Hanson et al., 2008; Landmesser, 2018; Shwartz et al., 2013). Spinal SNA starts at midgestation stages - around the 12^th^ day of embryonic development in mice (E12.5) - and is generated autonomously by the ventral spinal cord, before the establishment of sensory inputs from the periphery and descending inputs from the brain. Therefore, SNA is fully preserved in the isolated fetal spinal cord (Czarnecki et al., 2014; Hanson and Landmesser, 2003). Although its underlying molecular and cellular mechanisms have yet to be fully resolved, spinal SNA has been shown to involve spontaneous waves of electrical activity depolarizing cholinergic motoneurons as well as GABAergic and glutamatergic interneurons along the length of the ventral spinal cord (Czarnecki et al., 2014; Hanson and Landmesser, 2003; Hanson et al., 2008). By contrast, the electrophysiological behavior of ventral NEPs has never been characterized, despite report that they can exhibit calcium transients during episodes of SNA (Wang et al., 2009).

## RESULTS

To address this question, we first examined the electrical activity of NEPs using patch-clamp recordings on a preparation of whole spinal cord from E12.5 mouse embryos in the open-book configuration (Czarnecki et al., 2014; Osterstock et al., 2018). This preparation gave us access to all NEPs lining the central canal while preserving SNA generation (**Figure 1a)**. Five distinct bilateral domains of NEPs are classically identified along the central canal of the ventral spinal cord: p0, p1, p2, pMN and p3 domains (**Figure 1a)**. A sixth unique NEP domain called the floor-plate is located at the base of the spinal cord, joining both sides of the neuroepithelium at the ventral midline. Patch-clamp recordings first revealed that ventral NEPs exhibit spontaneous and recurrent depolarizations (**Figure 1b & c**) at an interval of 183 ± 38 s (N = 32 cells), similar to the interval previously reported in cholinergic motoneurons during spinal SNA (Czarnecki et al., 2014). Strikingly, the amplitude of NEP spontaneous depolarization was not uniform but was inversely correlated to the distance of the recorded NEPs from the ventral midline (**Figure 1c & d)**, the largest depolarization being observed in floor-plate NEPs. To determine whether NEP depolarizations were correlated between these different regions, we performed paired recordings between the floor-plate and the neighboring p3 domain (Distance _[FP-p3]_ = 22 ± 7 μm, N = 6; **Figure 1f, g & h)**. Simultaneous recordings demonstrated that depolarizations were indeed correlated between NEPs, although they exhibited significantly smaller amplitudes and a slight delay in the p3 domain when compared to the floor-plate NEP (**Figure 1g**). These results suggested that these depolarizing events propagated from the floor-plate to other NEP domains through direct electrical coupling. Indeed, neuroepithelial progenitors are known to be extensively coupled by gap-junctions at this developmental stage (Bittman et al., 2004). Confirming this observation, we found that neurobiotin filling via the patch pipette often labeled several adjacent cells in the p3 domain (14.7 ± 6.3 cells in p3, N = 6) as well as in the floor-plate (6.6 ± 2.9 cells in FP, N = 10) (**Figure 1f and 2d**). We also demonstrated that ventral NEPs formed an electrical syncytium by injecting current steps in NEPs located in the p3 domain. These current steps were systematically associated with a proportional, albeit much smaller, depolarization in floor-plate NEPs (Ratio _[FP/p3]_ = 4.6 ± 1.7 %, N = 18), showing that they are electrically coupled (**Figure 3c**). Taken together, our results indicated that NEP depolarizations observed during episodes of SNA originate from the floor-plate and propagate dorsally into the rest of the neuroepithelium through gap-junctions.

**Figure 1.**
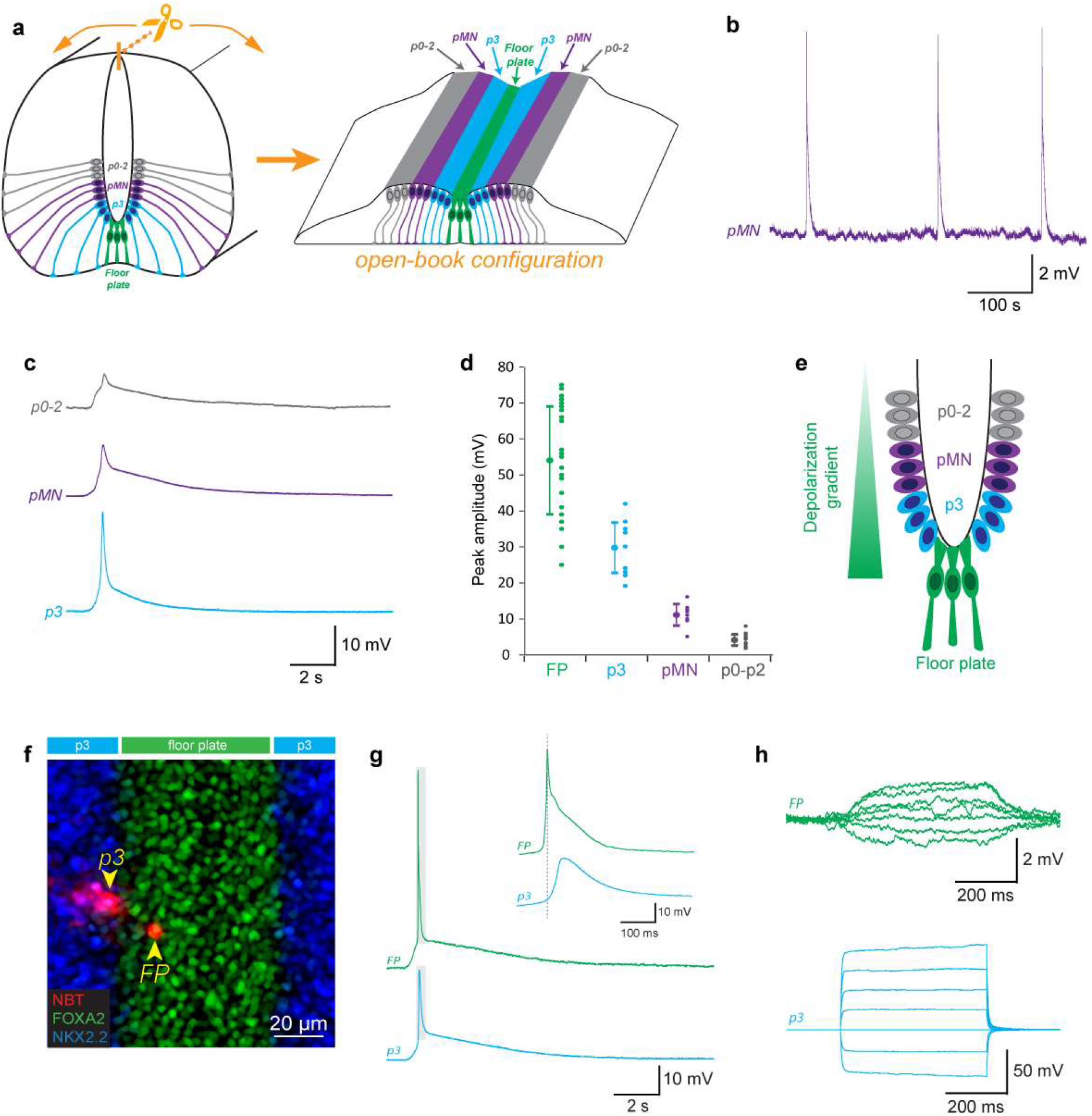
Neuroepithelial progenitors are depolarized during SNA following a ventro-dorsal gradient. **a,** Drawings showing the location and radial organization of the ventral neuroepithelial progenitor domains (p2-p0, pMN, p3, floor-plate) in a coronal section of an E12.5 mouse spinal cord (left) and in in the open-book configuration (right) used for patch-clamp recordings and calcium imaging. Domains were determined by their distance from the midline in open-book configuration (p3 = 30-70 μm; pMN = 70-110 μm; p0-p2 = 110-200). **b,** Example of current-clamp recording showing recurrent depolarization at the same frequency as spinal SNA in a neuroepithelial progenitor located in the pMN domain. **c,** Example of current-clamp recording showing spontaneous depolarizations recorded separately in neuroepithelial progenitors from the p0-p2 domain (top trace, in grey), the pMN domain (middle trace, purple) and the p3 domain (bottom trace, blue). **d,** Plot showing the average amplitude and amplitude distribution of spontaneous events recorded in different neuro-epithelial progenitor domains. Note that amplitudes are inversely correlated to the distance of the recorded domain from the midline. **e,** Drawing illustrating the depolarization gradient observed in the ventral neuroepithelium. **f,** Confocal image showing a pair of floor-plate and p3 neuroepithelial progenitors simultaneously recorded by dual patch-clamp and filled with neurobiotin (NBT, red). The recorded open-book preparation was processed for immunostaining against the transcription factor FoxA2 (floor-plate marker, green) and NKX2.2 (p3 domain marker, blue). The two recorded cells are indicated by yellow arrowheads. Note that several cells were filled by neurobiotin, indicating gap junction coupling between neuroepithelial progenitors. **g,** Example of a spontaneous depolarization simultaneously recorded in a floor-plate and a p3 neuro-epithelial progenitor. Note the reduced amplitude and slight delay of the p3 depolarization that is consistent with passive propagation of the depolarization from the floor-plate to the p3 domain through gap junctions. **h,** Injection of a depolarizing current in a p3 neuro-epithelial progenitor resulted in the simultaneous depolarization of a floor-plate cell, confirming the existence of functional gap junctions between these cells.

**Figure 2.**
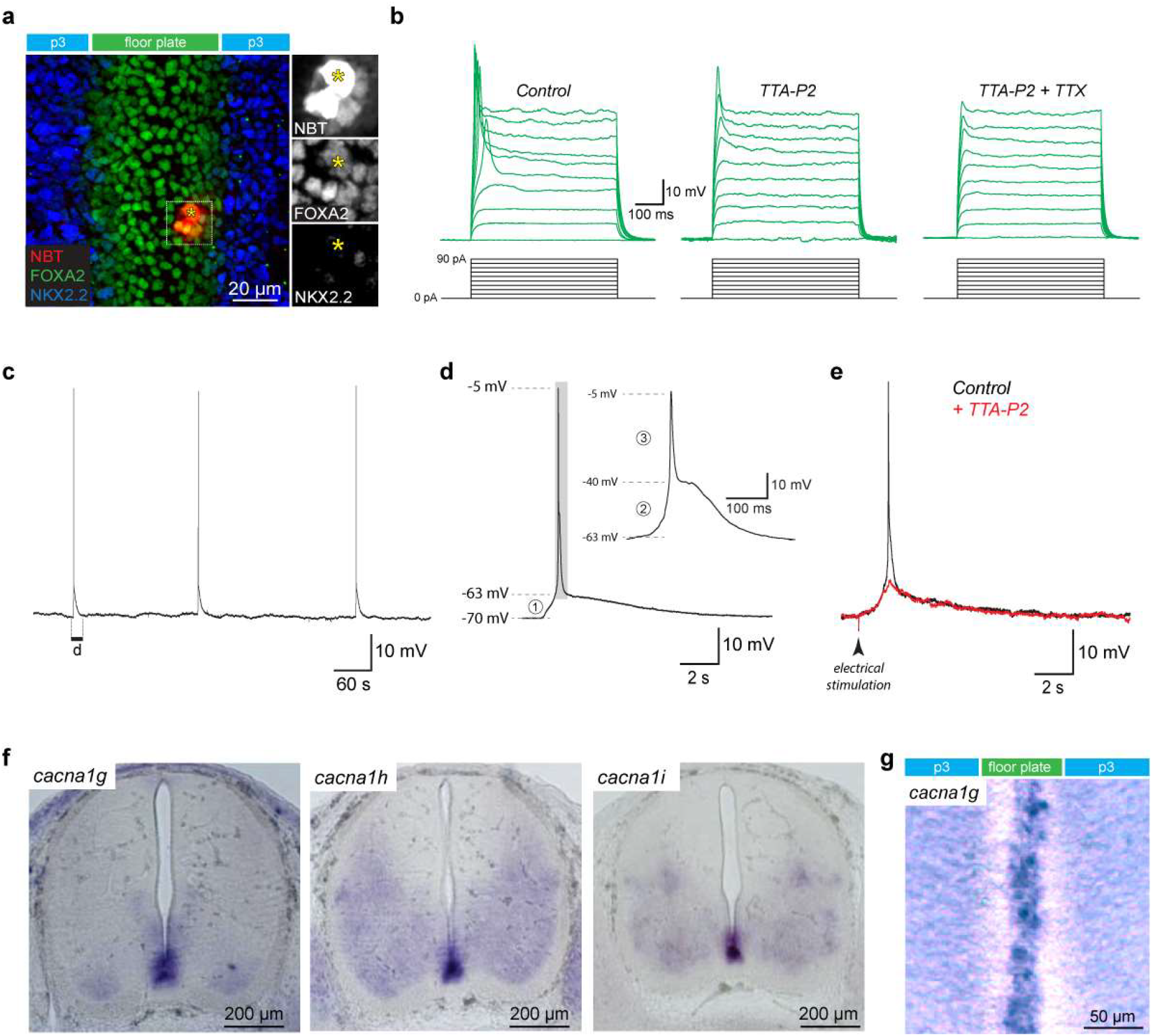
Floor-plate neuroepithelial progenitors generate recurrent action potentials during spinal SNA. **a,** Confocal image from an E12.5 spinal cord in open-book configuration after patch-clamp recording of a floor-plate cell. The recorded floor-plate cell (yellow asterisk) was filled with neurobiotin (NBT, red) and the spinal cord was processed for immunostaining against the transcription factor FoxA2 (green), expressed in floor-plate cells and against the transcription factor NKX2.2 (blue) expressed in neuroepithelial progenitors in the adjacent p3 domain. Note that several cells were filled by neurobiotin, indicating gap junction coupling between floor-plate cells. **b,** Example of current-clamp recordings obtained in a floor-plate cell in response to 10 incremental current steps (increment = 10pA) in control conditions (left traces), after application of the T-type calcium channel blocker TTA-P2 (3 μM; center traces) and subsequent addition of the sodium channel blocker TTX (1 μM; right traces). Note that action potentials triggered in floor-plate cells were only fully blocked by a combination of both channel blockers. **c,** Example of current-clamp recording from a floor-plate cell generating recurrent depolarization. **d,** Magnified view of a spontaneous depolarization revealing the presence of three distinct components: a small initial depolarization (1, from −70 to −63 mV) and a biphasic action potential (grey area magnified in insert) composed of a slow (2, from −63 to −40 mV) and fast component (3, from −40 to −5 mV). **e,** depolarization with a time course identical to spontaneous events could also be evoked by an electrical stimulation located 3 mm rostrally to the site of recording. Application of TTA-P2 (3 μM) fully blocked the biphasic action potential while the small initial depolarization remained unaffected. **f,***In situ* hybridization showing the location of mRNA transcripts for the 3 known sub-types of T-type channels (*cacna1g, cacna1h, cacna1i*) in coronal sections of E12.5 mouse embryos. Note that transcripts for all 3 subunits are highly expressed in the floor-plate. **g,***In situ* hybridization showing location of mRNA transcripts for *cacna1g* in an E12.5 spinal cord in open-book configuration.

**Figure 3.**
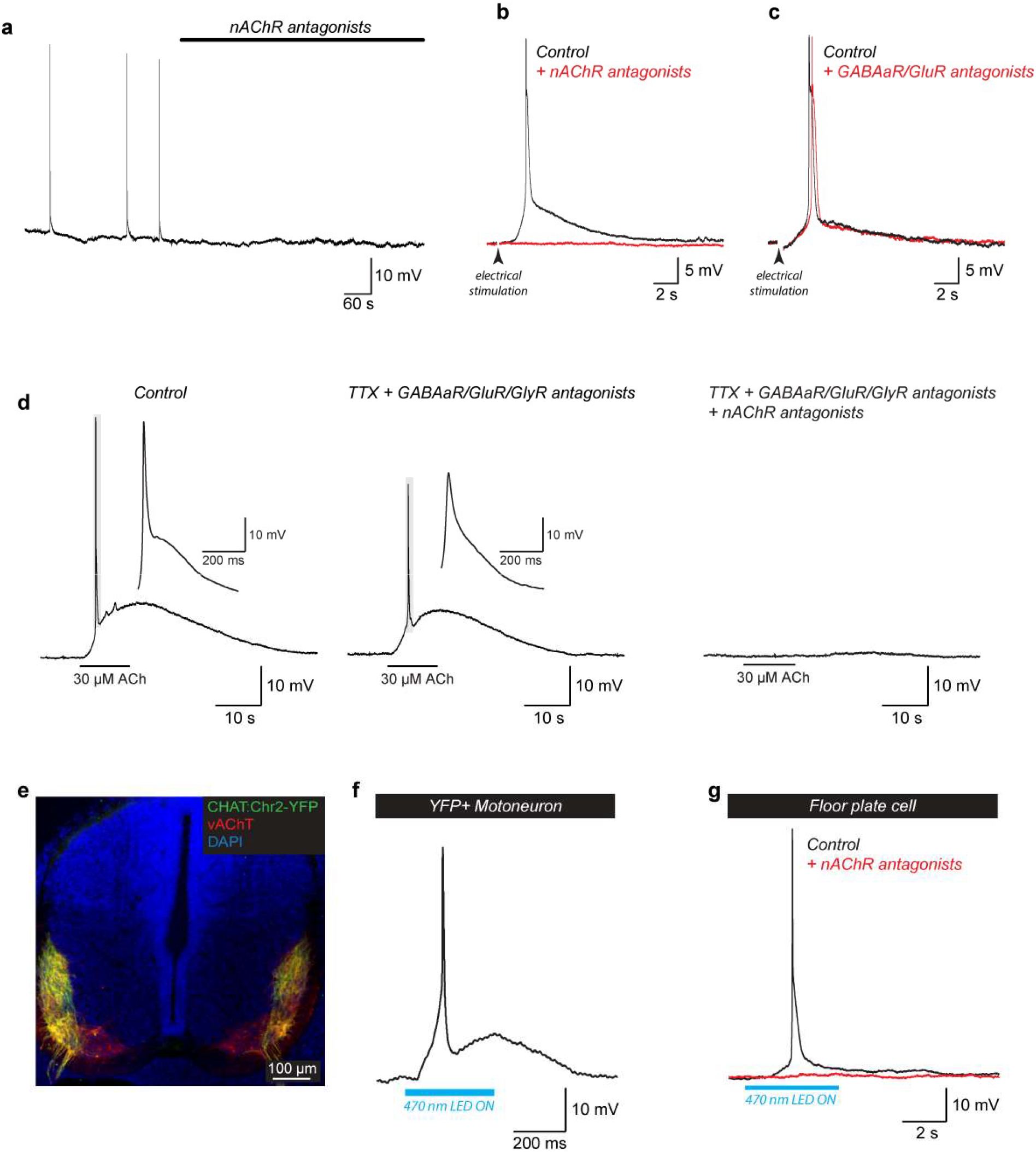
Floor-plate biphasic action potentials are triggered by the activation of nicotinic acetylcholine receptors in response to acetylcholine released by motoneurons. **a,** Example of recurrent spontaneous floor-plate action potentials blocked after addition of the nicotinic acetylcholine receptor (nAChR) antagonists: Mecamylamine (50 μM) and d-Tubocurarine (5 μM). **b,** Example of current-clamp recording showing floor-plate action potential evoked by electrical stimulation in control condition (black trace) and after addition of the nAChR antagonists (red trace). **c,** Example of current-clamp recording showing floor-plate action potential evoked in control condition (black trace) and after addition of antagonists against ionotropic receptor for GABA (Gabazine 3 μM) and glutamate (DL-APV 200 μM and CNQX 20 μM). **d,** Example of current-clamp recording showing that floor-plate action potential can be evoked by local application of 30 μM acetylcholine (left trace), even in the presence of TTX (1 μM) and antagonists to AMPA/Kainate glutamate receptor (CNQX 10 μM), NMDA glutamate receptor (DL-APV 200 μM), GABA_A_ receptor (Gabazine 3 μM) and glycine receptor (strychnine 1 μM) (center trace). Floor-plate action potential evoked by acetylcholine were blocked by the addition of nAChR antagonists (right trace). Note that the addition of TTX inhibited the fast component of the biphasic action potential (see **Supplementary Figure 2**). **e,** Confocal image of a coronal section from a ChAT:ChR2-YFP mouse embryo at E12.5 showing the expression of Channelrhodopsin2-YFP fusion protein (in green) in cholinergic motoneurons located in the ventro-dorsal horns and labelled with the vesicular acetylcholine transporter vAChT (in red). All cell nuclei were labelled using DAPI (in blue). **f,** Example of current-clamp recording from a ChAT:ChR2-YFP^+^ motoneuron showing an action potential triggered by the opening of Channelrhodopsin 2 in response to blue light stimulation (470 nm). **g,** Example of current-clamp recording from a floor-plate cell recorded in a ChAT:ChR2-YFP^+^ fetal spinal cord showing how blue light stimulation could evoke a slow cholinergic depolarization and trigger a biphasic action potential that were blocked by the addition of nAChR antagonists.

To understand how floor-plate NEPs could exhibit such large and recurrent depolarizations, we investigated their electrophysiological properties in more detail. In open-book configuration, floor-plate cells are easily identified by their location at the midline, their thin apical processes and their higher translucency when compared to surrounding neuroepithelial progenitor domains. To confirm the validity of these morphological criteria, some of the recorded cells were filled with the neurobiotin tracer and their molecular identity assessed by post-hoc immunostaining against the transcription factors FOXA2 and NKX2.2, which are respectively expressed in floor-plate NEPs and in the adjacent p3 neuroepithelial progenitor cells (Lupo et al., 2006). All floor-plate cells filled by neurobiotin were found to be FoxA2^+^ and NKX2.2^−^ (N= 10/10 cells; **Figure 1f and 2a**). Unlike other NEPs that sequentially generate neurons and glial cells during embryonic development (Rowitch, 2004), floor-plate NEPs display glial features at an early embryonic stage and have only been shown to generate ependymocytes – a subpopulation of glial cells lining the central canal of the adult spinal cord (Barry and McDermott, 2005; Cañizares et al., 2019; Deneen et al., 2006; Khazanov et al., 2017; Mirzadeh et al., 2017). Indeed, we found that all FoxA2^+^/NKX2.2^−^ cells co-expressed the glutamate astrocyte transporter GLAST, a marker for glial progenitors (**Supplementary Figure 1a-h**) together with the transcription factor Sox2, a marker for neuroepithelial progenitors (**Supplementary Figure 1i-l)**. Recorded floor-plate NEPs had an average membrane resistance of 480 ± 410 MΩ, an average membrane capacitance of 18.3 ± 6.6 pF and an average membrane potential of −64.8 ± 9.7 mV (N= 24 cells). More importantly, we found that floor-plate NEPs had the unique ability to generate action potentials (AP) in response to membrane depolarization (threshold = −38 ± 11 mV; amplitude = 44 ± 17 mV; N= 17 cells; **Figure 2b, left trace**). Unlike neuronal APs which only rely on voltage-gated sodium channels, floor-plate APs could only be blocked by a combination of TTA-P2 (**Figure 2b, middle trace**), a specific T-type calcium channel blocker, and TTX, a sodium channel blocker (**Figure 2b, right trace**; N = 15 cells). Voltage-dependent inward currents with T-type-like properties had been reported in cells located in the rat floor-plate at later developmental stages (Frischknecht and Randall, 1998). By contrast, NEPs recorded in the adjacent p3 domain did not exhibit any voltage-dependent responses (N= 24 cells; **Figure 1h**). During SNA, floor-plate recurrent depolarizations (amplitude = 54 ± 15 mV; N = 22 cells; **Figure 2c**) exhibited a complex and stereotypical time course. These spontaneous events were comprised of a small initial depolarization (amplitude = 6.9 ± 1.0 mV; half-width = 2.8 ± 1.0 s) followed by an AP exhibiting a biphasic time course comprised first of a slow component (amplitude = 18.9 ± 5.5 mV; half-width = 274 ± 57 ms) followed by a fast component (amplitude = 42.4 ± 7.2 mV; half-width = 11.8 ± 3.2 ms; **Figure 2d**). Identical events could be evoked using an electrical stimulation (32 mA, 200 μs) placed on the midline, 2 mm rostrally to the site of recording (**Figure 2e)**. Application of the T-type calcium channel blocker TTA-P2 (3 μM) was sufficient to fully block the biphasic action potential, while the small initial depolarization remained unaffected (N=8/8 cells). To confirm and further characterize the expression of T-type calcium channels in the floor-plate at this developmental stage, we performed *in situ* hybridization for the 3 known T-type calcium channels (*cacna1g*, *cacna1h, cacna1i*). We found that mRNA coding for T-type calcium channels were all highly and predominantly expressed in the floor-plate (**Figure 2f & 2g**).

Since SNA is dependent on acetylcholine released from motoneurons (Czarnecki et al., 2014; Hanson and Landmesser, 2003), we investigated whether recurrent floor-plate action potentials were triggered by this neurotransmitter. First, we found that both spontaneous and evoked floor-plate events were fully blocked by antagonists against nicotinic acetylcholine receptors (nAChRs; Mecamylamine and d-Tubocurarine; N = 12/12 cells; **Figure 3a & b**). By contrast, they remained unaffected by antagonists against GABA_A_ receptors (Gabazine) and glutamate ionotropic receptors (DL-APV and CNQX), two other neurotransmitters released by spinal interneurons during SNA (Czarnecki et al., 2014; Hanson and Landmesser, 2003) (**Figure 3c;** N = 11/11 cells). Exogenous application of acetylcholine (30 μM) was sufficient to depolarize floor-plate cells (Amplitude = 10.0 ± 1.5 mV) and trigger biphasic action potentials (amplitude = 59 ± 7 mV) in a nAChR dependent manner (N=6/6 cells) (**Figure 3d**). Application of the sodium channel blocker TTX together with antagonists of GABA, glutamate and glycine ionotropic receptors had no effect on the depolarization induced by acetylcholine, suggesting that acetylcholine acts directly onto nAChR expressed by floor-plate cells (**Figure 3d)**. We observed that TTX suppressed the fast component of the biphasic action potential triggered by acetylcholine, demonstrating that it resulted from the activation of voltage-gated sodium channels in floor-plate cells, while the slow component was blocked by TTA-P2 indicating that this resulted from the activation of T-type calcium channels (**Supplementary Figure 2**). To further confirm that ACh release from motoneurons triggers floor-plate action potentials, we performed optogenetic stimulation of cholinergic spinal motoneurons using a transgenic mouse model (ChAT:ChR2-YFP) where channelrhodopsin 2 is expressed in cholinergic motoneurons (**Figure 3e-f**). Using this approach, we confirmed that the release of endogenous acetylcholine triggered by optogenetic stimulation of motoneurons was sufficient to depolarize and trigger an action potential in floor-plate cells (N = 9 cells), which could be fully blocked by nAChR antagonists (N = 4/4 cells; **Figure 3g)**. Taken together, our results show that floor-plate action potentials are triggered by the activation of nAChR in response to acetylcholine released by motoneurons.

Finally, we examined whether and how floor-plate activity could also propagate between floor-plate cells along the rostro-caudal axis of the spinal cord. To investigate this possibility, we expressed the genetically encoded calcium indicator GCaMP6f in floor-plate cells using a conditional transgenic approach (GCaMP6floxP x GLASTCreERT2; see Methods). This was achieved by taking advantage of the fact that floor-plate cells are the first neuroepithelial cells to express the glutamate aspartate transporter (GLAST) during spinal cord development (Barry and McDermott, 2005; Deneen et al., 2006; Shibata et al., 1997) (**Figure 4a** and **Supplementary Figure 1**). Using this approach, we found that midline floor-plate cells expressing GCaMP6f (**Figure 4a**) generated spontaneous calcium waves at an interval of 176 ± 67 s (N = 10 embryos; **Figure 4b & c** and **Supplementary video 1**), similar to the interval observed for spontaneous floor-plate action potentials. Spontaneous floor-plate calcium waves propagated along the length of the spinal cord and could propagate in either direction. However, rostral to caudal waves initiated in the cervical area were more frequent (67%, 41/61 waves, N = 10 embryos) than caudal to rostral ones (33%, 20/61 waves) **(Supplementary Figure 3** and **Supplementary video 2)**. This proportion was similar to that previously reported for waves of muscle contractions observed *in vivo* in mouse embryos at the same developmental stage (Suzue and Shinoda, 1999). Average propagation speed of spontaneous rostro-caudal (2.4 ± 0.9 mm/s) and caudo-rostral floor-plate calcium waves (2.1 ± 1.3 mm/s) were not significantly different (paired Wilcoxon test, p=0.12, N = 10 embryos) and their speed was consistent with the propagation speed estimated for SNA waves in motoneurons (Hanson and Landmesser, 2003; Scain et al., 2010).

**Figure 4.**
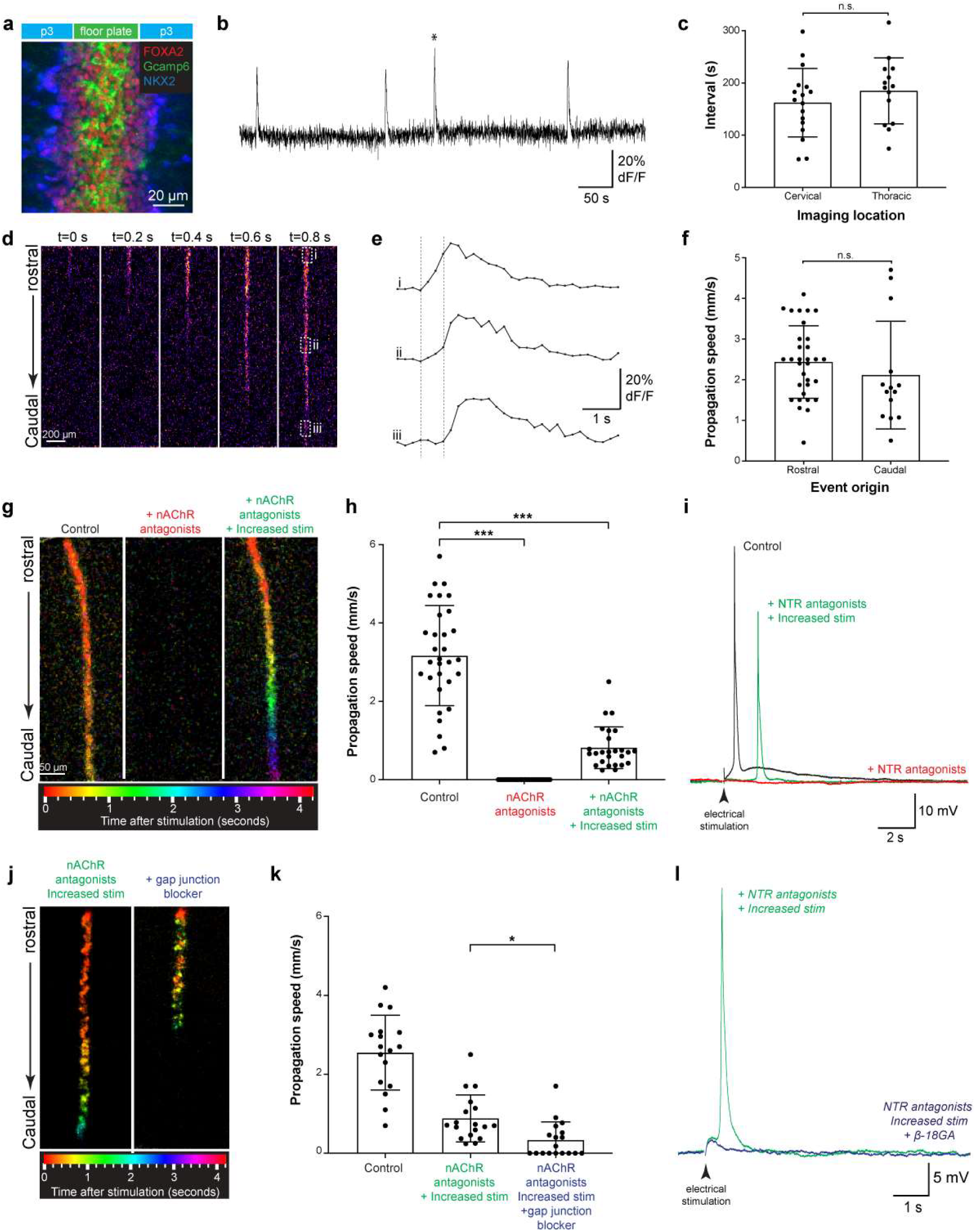
Floor-plate action potentials are associated with calcium waves and propagate along the length of the spinal cord. **a,** Confocal image showing the expression of the fluorescent GCaMP6 protein (in green) in floor-plate cells labelled with an antibody against FoxA2 (in red). Recombination of GCaMP6 was not induced in the p3 domain labelled with an antibody against NKX2.2 (in blue). **b,** Plot representing the intensity of fluorescence (dF/F) in the floor-plate over 10 min. Recurrent spontaneous calcium events were observed. Asterisk indicates the event represented in **d**. **c,** Plot quantifying the interval between spontaneous calcium events observed at cervical and thoracic segments. **d,** Time-lapse images showing the rostro-caudal propagation of GCaMP6 fluorescent signals in floor-plate cells over time in the thoracic region. Dotted boxes represent the region of interest used to calculate propagation speeds. **e**, Plot representing the intensity of fluorescence (dF/F) over time in the 3 regions of interest indicated in **d**. Dotted lines mark the onset of the calcium signals in region i (left) and iii (right). **f,** Plot of the propagation speed of floor-plate calcium waves according to the rostral or the caudal origin of the waves (N=61 waves; 10 embryos). No significant differences in propagation speed were observed between rostral and caudal origin. **g**, Images of color-coded calcium waves evoked by rostral electrical stimulation (32 mA, 200 μs) in control condition (left), after application of nAChR antagonists (center) and subsequent increase of stimulation intensity (32 mA, 1200 μs). Time after stimulation is coded in distinct color as indicated in the scale at the bottom. Note that calcium waves evoked at higher stimulation intensity in the presence of nAChR antagonists propagate more slowly. **h**, Plots of the propagation speed of evoked floor-plate calcium waves in different conditions. **i,** Example of current-clamp recording of a floor-plate cell in control condition (black trace), after application of a cocktail of antagonist to nAChR, GABA_A_R, iGluR and GlyR (red trace) and subsequent increase of stimulation intensity. Note that action potential could still be evoked and propagate even when nAChRs, GABA_A_Rs, iGluRs and GlyRs were blocked. **j,** Images of color-coded calcium waves evoked by increased electrical stimulation (32 mA, 1200 μs) in the presence of nAChR antagonists (left panel) and after addition of the gap junction blocker β18-GA (right panel). **k**, Plot of the propagation speed of evoked floor-plate calcium waves before and after addition of nAChR antagonists and gap junction blockers. **l**, Example of current-clamp recording of a floor-plate cell in the presence of a cocktail of antagonists to nAChR, GABA_A_R, iGluR and GlyR (green trace) and after subsequent addition of the gap junction blocker β18-GA (blue trace).

As observed for floor-plate action potentials, calcium waves could also be consistently triggered using electrical stimulation (32 mA, 200 μs, **Figure 4g, h, j, k** and **Supplementary Video 3**). Application of the T-type calcium channel blocker TTA-P2 (3 μM) was sufficient to fully block floor-plate calcium waves evoked by electrical stimulation (paired Wilcoxon test, p<0.001, N = 5 embryos; **Supplementary Video 4**), even after increasing the duration of stimulation (32 mA, 1200-1600 μs). The initiation and propagation of floor-plate calcium waves evoked by electrical stimulation was also blocked by the application of nAChR antagonists (paired Wilcoxon test, p<0.001, N = 6 embryos; **Figure 4g, Supplementary video 5**), as observed with floor-plate action potentials (**Figure 2**). However, increasing stimulation duration (32 mA, 1200-1600 μs) allowed the recovery of floor-plate calcium waves, albeit with a significantly decreased propagation speed of 0.8 ± 0.5 mm/s (paired Wilcoxon test, p<0.001, N = 6 embryos; **Figure 4g & h, Supplementary video 5**). Similarly, patch-clamp experiments showed that after increasing the duration of stimulation in the presence of a cocktail of antagonists to nAChRs, GABAaR, GluRs and GlyR, the electrical stimulus could directly trigger a delayed biphasic AP without the small initial cholinergic depolarization (N = 4 embryos; **Figure 4i**). This result shows that, although the initiation and subsequent propagation of floor-plate APs and calcium waves strongly depend on cholinergic transmission from motoneurons, direct electrical coupling between floor-plate cells (**Figure 1e**) also likely participates to this propagation. We tested this hypothesis by applying the gap-junction blocker β18-GA (Rozental et al., 2001). We found that it either reduced or blocked the propagation of the floor-plate calcium waves (paired Wilcoxon test, p<0.05, N=6 embryos; **Figure 4j & k, Supplementary video 5**) and action potentials (N=4 embryos; **Figure 4l**) that were evoked in the presence of nAChR antagonists and other neurotransmitter antagonists. Therefore, these results show that the floor-plate of the neuroepithelium possess the intrinsic ability to both generate and propagate electrical signals along the embryonic spinal cord.

## DISCUSSION

To our knowledge, the neuroepithelial floor-plate is the first example of a non-neuronal structure able to spontaneously generate and propagate action potentials in the developing central nervous system. Until now, these abilities were considered to be exclusive features of differentiating neurons. Accordingly, the ability to generate action potentials has been routinely used as a functional assay to confirm the neuronal phenotype of cells during development. In this context, although Ca^++^/Na^+^ biphasic action potentials relying on T-type calcium channels are not commonly observed in neurons, their previous discovery in neural cells in the Xenopus embryonic spinal cord (Gu et al., 1994) and in the mouse fetal hindbrain (Moruzzi et al., 2009) has been interpreted as an early and transient form of excitability in developing neurons. Instead, we found here that Ca^++^/Na^+^ biphasic action potentials are generated in the mouse spinal cord by gliogenic neuroepithelial progenitors located in the floor-plate and that they propagate laterally into other neuroepithelial cells also engaged into a gliogenic program at this stage (Deneen et al., 2006).

From an evolutionary point of view, the discovery of an epithelial electrical conduction system in the vertebrate fetal CNS should not come as a complete surprise. Indeed, epithelial and syncytial conduction systems are commonly observed in non-bilaterian animals exhibiting either a “primitive” nervous system, like jellyfish and comb jellies, or in those lacking a nervous system like the glass sponges (Anderson, 1980; Leys et al., 2007; Mackie, 1970; Meech, 2015). It has been proposed that the modern neuron-based nervous systems of bilaterians using axonal/chemical synaptic transmission evolved from such epithelial conduction systems coupled electrically by gap-junctions (Castelfranco and Hartline, 2016). This evolution would also have been accompanied by a switch from slow voltage-gated calcium channel - inherited from our protist ancestors and still found in the modern paramecium (Eckert et al., 1972) - to fast voltage-gated sodium channels (Castelfranco and Hartline, 2016). Our discovery of a “primitive” electrical conduction system in the neuroepithelium that gives rise to the “modern” nervous system of vertebrates during fetal development could therefore be interpreted as strong evidence supporting this evolutionary scenario.

One may also notice the functional similitude existing between the floor-plate of His and the heart electrical conduction system known as the bundle of His. The bundle of His was discovered by the anatomist and cardiologist Wilhem His, Jr. in 1893 (BAST and GARDNER, 1949), five year after his father had described the floor-plate (Kingsbury, 1920). Both structures act as electrical conduction systems connected by gap junctions able to propagate electrical signals across their respective organs. By showing that these structures not only share the same name but also share similar functional properties, our work reveal that the nervous and cardiac physiology of vertebrates is more deeply related than previously thought.

Further work will be needed to decipher what function(s) are fulfilled by this non-neuronal electrical conduction system. Although the electrical excitability of floor-plate cells could only be a vestigial property inherited from non-bilaterian ancestors, it is more likely playing a significant role in the development of the spinal cord. Indeed, the energetic cost of generating and propagating electrical signals (Attwell and Laughlin, 2001) as well as the redundancy of T-type channel subtypes expressed in the floor-plate point towards having a conserved and significant function. One hypothesis is that floor-plate excitability regulates and coordinates the release of the morphogen Sonic Hedgehog (Shh), required for neural tube patterning, or the expression of axon guidance molecules such as Slit and Netrin. Indeed, recent studies have shown unexpected interactions between early calcium electrical excitability and Shh signalling in the *Xenopus* spinal cord (Belgacem et al., 2015; Borodinsky and Belgacem, 2016). Moreover, T-type channel mutants have recently been identified in a forward genetic screen for neural tube defects in *Ciona intestinalis* (Abdul-Wajid et al., 2015), suggesting that the electrical gradient established by the floor-plate in the ventral neuro-epithelium could represent a novel morphogenetic mechanism. An additional and non-exclusive hypothesis is that floor-plate action potentials initiated by acetylcholine released by motoneurons during SNA act as a feedback signal to promote the switch from neurogenesis to gliogenesis in ventral neuro-epithelial progenitors (Barry and McDermott, 2005; Deneen et al., 2006) since both events occur around the same developmental stage.

In conclusion, our discovery of the floor-plate of His acting as a non-neuronal electrical conduction system could profoundly change how we conceive the origin, extent and role of electrical signals during the development of the central nervous system of vertebrates.

## SUPPLEMENTARY FIGURES

**Supplementary Figure 1.**
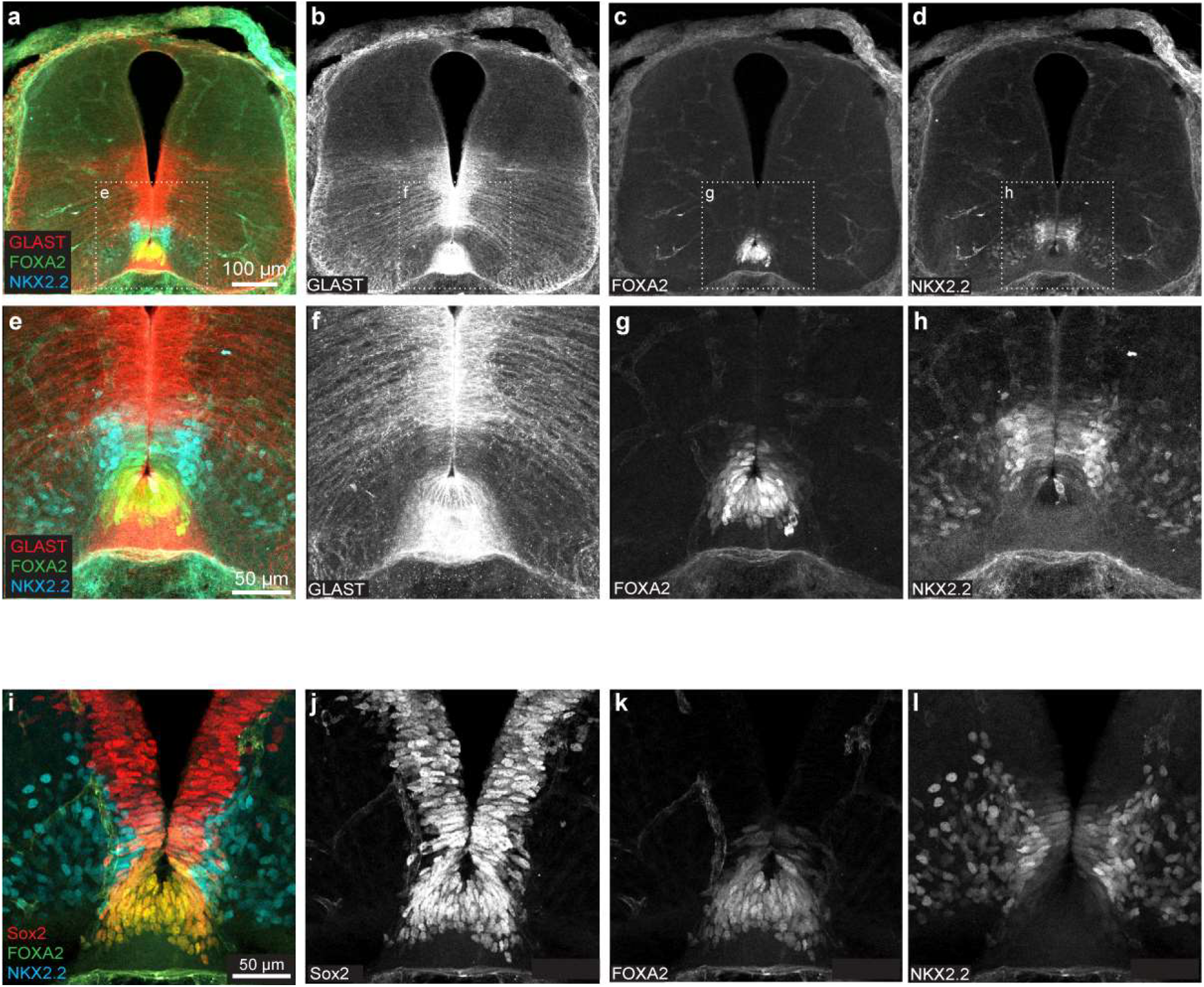
Floor-plate cells co-express GLAST, a marker of glial progenitors, and Sox2, a marker of neuro-epithelial progenitors. **a-d,** Confocal images showing the expression of the glial marker GLAST (red in a, greyscale in b), the floor-plate marker FoxA2 (green in a, greyscale in c) and the p3 domain marker Nkx2.2 (cyan in a, greyscale in d) in a coronal section of a E12.5 mouse embryonic spinal cord. **e-h** Higher magnification confocal image of the same coronal section shown in a-d (dotted area). Note that FoxA2^+^/NKX2.2^−^ cells located in the floor-plate co-expressed the glutamate astrocyte transporter GLAST. **i-l,** confocal image showing the expression of the neuro-epithelial marker Sox2 (red in i, greyscale in j), the floor-plate marker FoxA2 (green in i, greyscale in k) and the p3 domain marker Nkx2.2 (cyan in i, greyscale in l) in a coronal section of a E12.5 mouse embryonic spinal cord. Note that all FoxA2^+^/NKX2.2^−^ cells located in the floor-plate co-expressed Sox2.

**Supplementary Figure 2.**
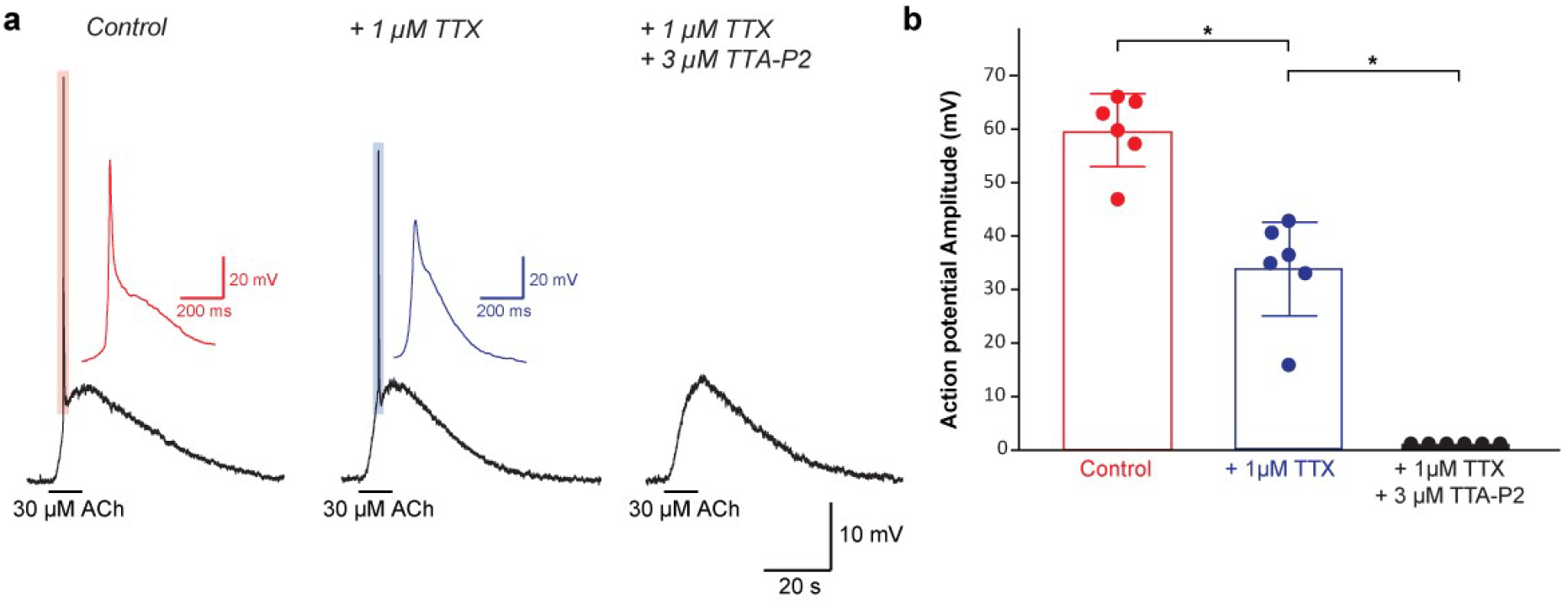
Floor-plate action potentials evoked by acetylcholine rely on both TTX-sensitive voltage-gated sodium channels and T-type voltage-gated calcium channels. **a**, Example of current-clamp recording showing a biphasic floor-plate action potential evoked by the application of acetylcholine (30 μM, left trace), after application of the voltage-gated sodium channel blocker TTX (1μM, middle trace) and after the subsequent addition of the T-type voltage-gated calcium channel blocker TTA-P2 (3 μM, right trace). Note that the fast component of the action potential was blocked by TTX while the slow component was only blocked after addition of TTA-P2. **b**, Plot quantifying the effect of TTX and TTA-P2 on the amplitude of floor-plate action potential evoked by acetylcholine.

**Supplementary Figure 3.**
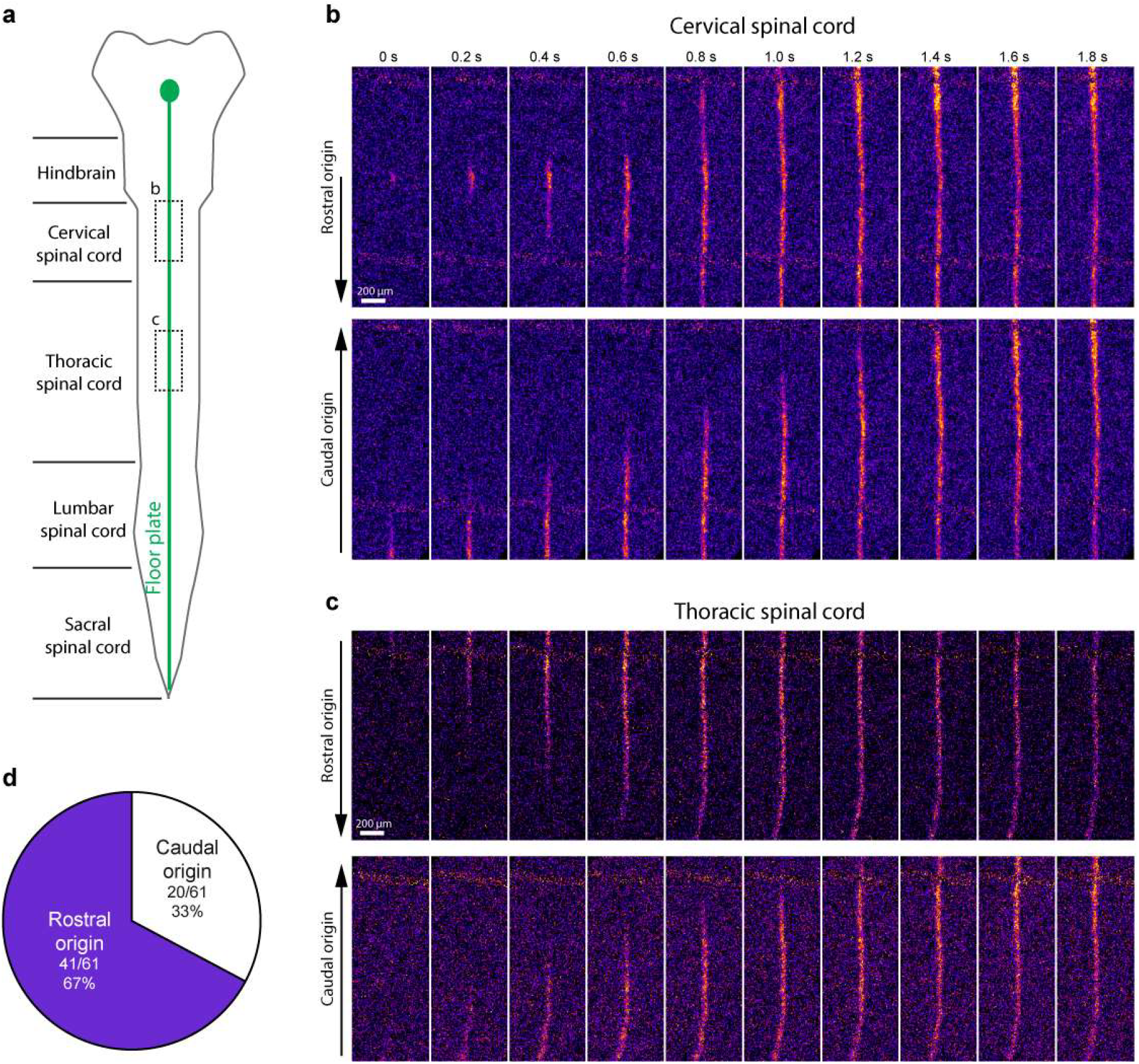
Spontaneous calcium waves in the floor-plate can propagate in both directions and originate from rostral or caudal locations. **a**, Drawing representing a longitudinal view of an open-book preparation of the spinal cord at E12.5. The floor-plate is represented in green. **b**, Time lapse images showing the evolution of GCaMP6 fluorescent signal in the floor-plate at the cervical level using an open-book preparation of spinal cord from a E12.5 transgenic mouse embryo (GLAST:CreERT2 x floxed- GCaMP6f). Examples of spontaneous floor-plate calcium waves with a rostral origin (top panel) and caudal origin (bottom panel) are shown. In the top traces, the wave originated from the cervical area and propagated in both directions. **c**, Time lapse images showing the evolution of GCaMP6 fluorescent signal in the floor-plate at the thoracic level. An example of each type of wave is shown (top panel: rostral origin, bottom panel: caudal origin). **d**, Proportion of spontaneous calcium wave originating respectively from a rostral and caudal origin in both regions (N = 61 waves; 10 embryos).

## SUPPLEMENTARY VIDEOS

**Supplementary Video 1. Example of background subtraction performed on a spontaneous calcium wave recorded at thoracic level.**

Left, Raw time-lapse recording (10 Hz) of GCaMP6 fluorescent signal showing a spontaneous floor-plate calcium wave recorded at thoracic level using an open-book preparation of spinal cord from a E12.5 transgenic mouse embryo (GLAST:CreERT2 x floxed-GCaMP6). Right, The same time-lapse recording (10 Hz) after subtraction of background GCaMP6 fluorescence.

**Supplementary Video 2. Spontaneous floor-plate calcium waves can propagate in either direction.**

Time lapse movies (10 Hz, background subtraction) showing two examples of floor-plate calcium waves recorded at thoracic level in the same open-book preparation exhibiting either a rostral (left movie) or a caudal origin (right movie).

**Supplementary Video 3. Evoked floor-plate calcium waves are consistently triggered by consecutive electrical stimulation.**

Time lapse movies (10 Hz, background subtraction) showing 3 consecutive floor-plate calcium waves recorded at thoracic level in response to electrical stimulation (32 mA/200 μs) located in the cervical area.

**Supplementary Video 4. Evoked floor-plate calcium waves are completely blocked by the T-type calcium blocker TTA-P2.**

Time lapse movies (10 Hz, background subtraction) showing examples of a calcium wave evoked in control condition (left movie; 32 mA/200 μs), after addition of TTA-P2 (middle movie; 32 mA/200 μs) and after subsequent increase of stimulation duration (right movie; 32 mA/1200 μs). Note that the T-type calcium channel blocker TTA-P2 (3 μM) completely blocked the floor-plate calcium wave, which is not restored after increasing the duration of electrical stimulation.

**Supplementary Video 5. Evoked floor-plate calcium wave propagation depends both on nAChR and gap-junction coupling.**

Time lapse movies (10 Hz, background subtraction) showing examples of a calcium wave evoked in control condition (first movie; 32 mA/200 μs), after addition of nAChR antagonist (second movie; 32 mA/200 μs), after subsequent increase of stimulation intensity (third movie; 32 mA/1200 μs) and after final addition of the gap-junction blocker β18-GA (fourth movie). Note that the floor-plate calcium wave propagation was partially restored by increasing the intensity of electrical stimulation and that it was subsequently reduced by addition of gap junction blocker.

## ACKNOWLEDGEMENTS

We thank Isabelle Dusart, Marine Robouam, Odile Favière, Goran Fort and Maryse Dardenne for their help in managing mouse colonies; Susanne Bolte, Jean-François Gilles and France Lam for assistance with confocal imaging. We thank Frank Pfrieger for the GLASTCreERT2 mouse line. This study was financially supported by Institut de Recherche sur la Moelle et l’encéphale (JMM), AFM-Telethon (JMM), Agence Nationale de la Recherche (JMM) and Fondation pour la Recherche Médicale (JMM and PL).

## AUTHOR CONTRIBUTIONS

K.H. Arulkandarajah, G. Osterstock and H. Le Corronc designed, performed and analysed the electrophysiological recordings. K.H. Arulkandarajah, G. Osterstock, S. Corsini and B. Le Bras designed, performed and analysed the immunohistochemistry experiments. A. Lafont, K.H. Arulkandarajah and E. Hong designed, performed and analysed calcium imaging experiments. K.H. Arulkandarajah, J. Boeri and A. Czarnecki designed, performed and analysed optogenetic experiments. N. Escalas, C. Soula, K.H. Arulkandarajah and E. Hong designed, performed and analysed *in situ* hybridization experiments. C. Mouffle and E. Bullier managed the animal colony, performed animal genotyping and contributed to immunohistochemistry experiments. J-M Mangin participated to the design and analysis of the experiments, supervised the project and wrote the manuscript with P. Legendre, C. Soula and E. Hong.

## COMPETING INTERESTS

The authors declare no competing interests.

## DATA AVAILABILITY

The authors declare that the data supporting the findings of this study are available within the paper.

## MATERIAL AND METHODS

### Mouse strains and transgenic models

All experiments were performed in accordance with European Community guiding principles on the care and use of animals (86/609/CEE, CE Off J no. L358, 18 December, 1986), French decree no. 97/748 of October 19, 1987 (J Off République Française, 20 October, 1987, pp. 12245–12248) and recommendations from the CNRS. All efforts were made to minimize the suffering of animals. In order to maximize litter size, experiments were performed on E12.5 mouse embryos obtained by crossing C57BL/6 males with females from a Swiss strain. For optogenetic stimulation of cholinergic motoneurons, we used C57BL/6 breeding male mice expressing a BAC transgene containing a Channelrhodopsin2-EYFP sequence driven by a Choline Acetyl Transferase (ChAT) promoter (ChAT-ChR2–EYFP, The Jackson Laboratory, USA, https://www.jax.org/strain/014546)(Zhao et al., 2011). For calcium imaging of floor-plate cells, we used C57BL/6 breeding male mice expressing a genetically encoded floxed Calcium indicator GCaMP6f (Ai95, The Jackson Laboratory, USA, https://www.jax.org/strain/028865), crossed with Swiss female mice expressing a tamoxifen-inducible Cre recombinase under the control of the glutamate aspartate transporter promoter (GLASTCreERT2, gift from Dr Frank Pfrieger)(Slezak et al., 2007). Cre recombination was induced by oral gavage of pregnant mice with 5 mg of tamoxifen (Sigma-Aldrich, France, T-5648) at embryonic days E10.5 and E11.5(Eon et al., 2008). Tamoxifen was dissolved in corn oil at 37°C for several hours to prepare 20 mg/mL solution(Zervas et al., 2004).

### In situ RNA hybridization staining

Mouse embryos were fixed in 3.7% paraformaldehyde (PFA) in PBS overnight at 4°C. Tissues were then sectioned at 60–80 μm using a vibratome (Microm, France). In all experiments, sections were performed at the cervico/brachial level. In situ hybridization (ISH) was performed on vibratome sections automatically (InsituPro, Intavis, Germany) using the whole-mount ISH as previously reported(Touahri et al., 2012). Digoxigenin (DIG)-labeled antisense RNA probes were synthesized using T3 polymerase from the following I.M.A.G.E. Consortium cDNA Clones(Lennon et al., 1996): *cacna1g*, (ID 6410519); *cacna1h* (ID 6820407) and *cacna1i* (ID 6825802). RNA labeled probes were detected by an alkaline-phosphatase coupled antibody (Roche Diagnostics), and NBT/BCIP (nitroblue tetrazolium/5-bromo-4-chloro-3-indolyl phosphate) were used as a chromogenic substrate for the alkaline phosphatase (Boehringer Mannheim, Germany). Stained sections were digitized and analyzed using Zeiss acquisition software (Zeiss, Germany).

### Isolated embryonic spinal cord preparation

E12.5 embryonic mouse spinal cords were dissected as previously described(Czarnecki et al., 2014; Osterstock et al., 2018). Briefly, pregnant mice were killed using a lethal dose of CO_2_ followed by cervical dislocation. Mice embryos of both sexes were dissected and whole spinal cord were left to recover for at least 30 min at 32°C in artificial cerebrospinal fluid (ACSF) containing 125 mM NaCl, 25mM NaHCO3, 11 mM Glucose, 3 mM KCl, 1 mM MgCl2, 2 mM CaCl2, and 1 mM NaH2PO4 (pH 7.3; 307 mOsm) continuously bubbled with 95% O_2_ – 5% CO_2_ gas, before being recorded or imaged. For ChAT-ChR2-EYFP and GCaMP6floxP x GLASTCreERT2 transgenic embryos, their tails were cut and observed under a fluorescent stereo-microscope (Leica MZ FLIII) to identify positive embryos for EYFP and GCaMP6, respectively.

### Patch-clamp recordings

Isolated spinal cords were placed in open-book configuration in a recording chamber under a nylon holding grid and continuously perfused (2 ml/min) at room temperature with the oxygenated ACSF described above. In contrast to previous studies(Czarnecki et al., 2014; Osterstock et al., 2018), the orientation of open-book spinal cord was inverted so that the neuro-epithelial layer was facing upward. Whole-cell voltage-clamp recordings of floor-plate cells and other neuroepithelial progenitors were performed under direct visualization using an infrared-sensitive CCD video camera. In addition to their location at the midline, floor-plate cells were distinguished from surrounding neuro-epithelial progenitors based on their smaller apical process and their higher translucency when using brightfield microscopy. To confirm their identity, some recorded cells were filled with neurobiotin (1 mg/ml, Vector labs) and revealed in combination with FoxA2 and Nkx2.2 immunostaining. Whole-cell patch-clamp electrodes were pulled from thick-wall borosilicate glass using a Brown-Flaming puller (P97, Sutter Instrument, USA). The tip of the electrode was fire-polished using a microforge (MF-830, Narishige, Japan). Patch-clamp electrodes had tip resistances comprised between 4 and 7 MΩ. Electrodes were filled with an intracellular solution containing either (in mM): 96.4 K^+^-methanesulfonate, 33.6 KCl, 4 MgCl2, 4 Na_2_ATP, 0.3 Na_3_GTP, 10 EGTA, and 10 HEPES or 100 K^+^-gluconate, 34 KCl, 0.5 EGTA, 10 HEPES, 4 MgCl_2_, 4 Na_2_ATP (pH 7.2; 290 mOsm). In both solutions, the theoretical equilibrium potential for chloride ions (ECl) was −30 mV, while cation reversal potential (Ecat) was 0 mV. Signals were recorded with a low-pass filtered at 4 kHz using a Multiclamp 700B amplifier (Molecular Devices, San Jose, CA, USA) and digitized at 20 kHz using a Digi-data 1440A interface coupled to pClamp 10.5 software (Molecular Devices, San Jose, CA, USA) running on a PC computer. Cells exhibiting a resting membrane potential above −50 mV (I=0 pA) were discarded from further analysis.

### Electrical and optogenetic stimulation

For electrical stimulations experiments, we used bipolar platinum electrodes (FHC, Bowdoin, ME, USA) placed 2 mm rostrally to the recording site onto the midline. For optogenetic stimulation of ChAT-ChR2 we used a 400 μm diameter optical fiber coupled with a 470 LED (Thorlabs, Germany) and centered onto the recording site. Electrical and optogenetic stimulations were performed at a low frequency (0.005 Hz, 1 stimulation every 180 s) in order to evoke a stable cholinergic response.

### Calcium imaging

Isolated spinal cords were placed in open-book configuration under a nylon holding grid in a recording chamber and continuously perfused (2mL/min) with the same oxygenated ACSF described above. Recombined fluorescent cells were observed under an epifluorescence microscope BX51W1 (Olympus) with a 470-515nm fluorescent excitation light to confirm GCaMP6f expression in the floor-plate. Time-lapse imaging was performed using either an Orca Flash4.0 (Hamamatsu, Japan) or Orca 03G (Hamamatsu, Japan) camera and recorded using the HC Image Live software (Hamamatsu, Japan) at an acquisition frequency of 5 or 10Hz with an exposure time of 200 ms and 100 ms, respectively. Images were encoded using a 256×256 or 512×512 pixel resolution and 16-bit grey scale. Time-lapse images were analyzed using the ImageJ software (National Institute of Health, USA). Signals were either extracted by subtracting each frame from the one following it (Figure 4g & 4i) or by subtracting an average of 10 frames preceding the onset of an event (Figure 4d, Supplementary Figure 3 and Supplementary videos 1-5). To measure the propagation speed, we pinpointed three regions of interest on the midline and measured the delays necessary to reach half-peak between each region of interest and averaged them. Intensity of signals (dF/F) were measured by dividing the peak amplitude of the events by the baseline value in raw images.

### Pharmacological agents

In most experiments, drugs were applied via the bath perfusion system. Typically, the effect of drugs reached equilibrium within 3 to 5 minutes. For acetylcholine application, we used a 0.5 mm diameter quartz tubing positioned 50 μm away from the recording area under direct visual control. Typically, response to acetylcholine reached equilibrium a few seconds (6 sec) after solution switch. The quartz tubing was connected using a manifold to six solenoid valves linked with six reservoirs. Solutions were gravity-fed into the quartz tubing. Drug application was controlled using a VC-8 valve controller (Warner Instruments, USA). At least 3 events were recorded in each condition after waiting 5 min for drugs to act. The following pharmacological agents were used: Acetylcholine chloride (30 μM, Sigma-Aldrich, Germany), TTX (1 μM, Alomone Labs, Israel), gabazine (3 μM, Tocris, USA), CNQX Disodium (20 μM, Tocris, USA), DL-APV (200 μM, Tocris, USA), Mecamylamine hydrochloride (100 μM, Tocris, USA), D-turbocurarine chloride (10 μM,Tocris Bioscience, Minneapolis, MN, USA), 18-β glycyrrhetinic acid (50 μM, Sigma-Aldrich, Germany) and TTA-P2 (3μM, Alomone, Israel). All drugs were dissolved at final concentration in ACSF.

### Immunohistochemistry

E12.5 embryos or whole embryonic spinal cord were immersion-fixed in phosphate buffer saline (PBS) with 4% PFA freshly prepared in PBS (pH 7.4) for 1 h at 4°C. Embryos were then rinsed with PBS and cryoprotected in PBS-15% sucrose at 4°C for 24 h and then in PBS-30% sucrose at 4°C for 24 h. Embryos were embedded in OCT medium (VWR, USA) and quickly frozen. Serial sections 20 μm thick were collected onto slides using a cryostat (Leica, Wetzlar, Germany) and stored at −80°C until immunohistochemical studies. For immunostaining, slides and whole spinal cord were washed in PBS, incubated in NH_4_Cl (50 mM) diluted in PBS for 20 min and then permeabilized for 30 min in a blocking solution (10% goat serum in PBS) with 0.2% Triton X-100. They were incubated overnight at 4°C with the primary antibodies, which were diluted in the 0.2% Triton X-100 blocking solution. Slides or whole spinal cords were then washed in PBS and incubated for 2h at room temperature in the appropriate secondary antibodies (diluted at 1/1000 in the 0.2% Triton X-100 blocking solution). The following primary antibodies were used: a mouse monoclonal anti-HNF-3β/FoxA2 (H-4) antibody sc-374376 (1:300, Santa Cruz Biotechnology, USA), a mouse monoclonal anti-Nkx2.2 antibody 74.5.A5 (1:600, Developmental Studies Hybridoma Bank, USA), a guinea-pig polyclonal anti-EAAT1/GLAST antibody 250 114 (1:1000, Synaptic System, Germany), a rabbit polyclonal anti-VAChT antibody (1:1000, Synaptic System, Germany) and a rabbit polyclonal anti-GFP antibody (1:1000, Invitrogen, USA). Alexa Fluor 405-, 488-, or 594- and 647-conjugated secondary antibodies (1/1000; Invitrogen, USA) were used to detect mouse monoclonal, guinea pig and rabbit polyclonal primary antibodies. After washing in PBS, slides or whole spinal cords were dried and mounted in Mowiol medium (Millipore, USA).

### Confocal microscopy

Once mounted, immunostained sections were imaged using a SP5 confocal microscope (Leica, Germany) using a 20X oil-immersion objective with a numerical aperture of 1.25, as well as with a 63X oil-immersion objective with a numerical aperture of 1.32 and a 1X digital zoom magnification. Serial optical sections were acquired with a Z-step of 3 μm (20X) and 0.78 μm (63X). Images (1024 × 1024; 8 to16 bit grayscale) were acquired using Leica software LAS-AF and analyzed using ImageJ (National Institutes of Health, USA). Colocalization with neurobiotin staining was assessed in three axes using a single confocal slice and x and y orthogonal views of the stack (ImageJ 1.5).

### Statistics

Statistical comparisons for electrophysiological recordings and calcium imaging parameters were conducted using GraphPad Prism 8 by performing non-parametric Wilcoxon-Mann-Whitney U tests for independent samples and Wilcoxon signed rank test for paired samples. We determined that statistical significance was obtained when the two-tailed p-value was <0.05.

## Notes

### Competing Interest Statement

The authors have declared no competing interest.

